# SCHNAPPs - Single Cell sHiNy APPlication(s)

**DOI:** 10.1101/2020.06.07.127274

**Authors:** Bernd Jagla, Vincent Rouilly, Michel Puceat, Milena Hasan

## Abstract

**Motivation:** Single-cell RNA-sequencing (scRNAseq) experiments are becoming a standard tool for bench-scientists to explore the cellular diversity present in all tissues. On one hand, the data produced by scRNASeq is technically complex, with analytical workflows that are still very much an active field of bioinformatics research, and on the other hand, a wealth of biological background knowledge is often needed to guide the investigation. Therefore, there is an increasing need to develop applications geared towards bench-scientists to help them abstract the technical challenges of the analysis, so that they can focus on the Science at play. It is also expected that such applications should support closer collaboration between bioinformaticians and bench-scientists by providing reproducible science tools.

**Results:** We present SCHNAPPs, a computer program designed to enable bench-scientists to autonomously explore and interpret single cell RNA-seq expression data and associated annotations. The Shiny-based application allows selecting genes and cells of interest, performing quality control, normalization, clustering, and differential expression analyses, applying standard workflows from Seurat (Stuart et al., 2019) or Scran (Lun et al., 2016) packages, and most of the common visualizations. An R-markdown report can be generated that tracks the modifications, and selected visualizations facilitating communication and reproducibility between bench-scientist and bioinformatician. The modular design of the tool allows to easily integrate new visualizations and analyses by bioinformaticians. We still recommend that a data analysis specialist oversees the analysis and interpretation.

**Availability:** The SCHNAPPs application, docker file, and documentation are available on GitHub: https://c3bi-pasteur-fr.github.io/UTechSCB-SCHNAPPs; Example contribution are available at the following GitHub site: https://github.com/baj12/SCHNAPPsContributions.

## Introduction

A successful and efficient analysis of data from single cell experiments requires a close interaction between bioinformaticians and bench-scientists. While the first feeds the data through an analysis pipeline, the second interprets the results. This is a challenging collaboration since the bioinformatician usually doesn’t have the biology-specific knowledge to select the cells and/or genes of interest and the bench-scientist may have difficulties handling the technical aspects of the analysis. It is common that a pipeline must be rerun multiple times to remove cells or genes from the analysis because they belong to cell types or to biological processes that are not relevant to the scientific question. For example, mitochondrial and ribosomal gene expression is separate from most biological processes and should be excluded during analysis. This implies numerous iterations to discuss intermediate results.

Many tools are being developed to tackle this challenge, some of which are covered in a recent review (Çakır et al. 2020). Among the most accomplished are iSEE (Rue-Albrecht et al. 2018), Cerebro (Hillje R.,et al. 2020), ASAP (Gardeux et al. 2017), iS-CellR (Patel MV 2018), and singleCellTK (Jenkins D, et al. 2019).

Here, we present SCHNAPPs (Single Cell sHiNy APPlication(s)), an R/Shiny application that has been designed to aid the communication between bench-scientists and bioinformaticians. Thus, many meetings can be avoided and the time for analyzing the data can be significantly reduced. The bench-scientist is enabled to characterize individual cells and genes starting from the initial normalization steps of raw counts to differential expression analysis. The selection process is captured in a report that can be used by a bioinformatician to validate and optimize the results. The software architecture of the application makes it easy for bioinformaticians to integrate new visualizations and analyses.

SCHNAPPs, has been successfully applied to validate a human “in a dish” model for a valvular disease (Neri T, et al, 2019) and to show that epicardium activation during a cardiomyopathy gives rise to both adipocytes and fibroblasts (Suffee N, et al., 2020). It is a standard tool for the analysis of scRNAseq data at our facility.

## Implementation

**Input** to the application is either a simple count matrix of comma-separated values (CSV), with rows representing features/genes and columns representing cells, or a SingleCellExperiment object (Lun A. and Risso D., 2019) with a sparse matrix holding the non-normalized counts and annotations for the cells (covariates) and annotation data for the features/genes. The singleCellExperiment object must have the following gene information for each gene: “symbol”, the gene-symbol; “id”, a potentially different unique identifier; “Description”, descriptive information for the gene. Cell specific information must include “sampleNames”, a string/factor to distinguish cells from different samples; and “barcode”, a unique barcode per sample. In practice, additional gene specific annotations like functional annotations from Ensembl (Yates et al. 2020), or cell specific annotations like cell type predictions using from SingleR (Aran D. et al., 2019) are computed by the bioinformatician during data preparation on the command line and integrated into the singleCellExperiment object.

Examples for generating these objects are given on GitHub (https://github.com/baj12/SCHNAPPsContributions#prepare-data-for-schnapps). Additional annotations for cells or genes can be loaded via CSV files. This way other types of information can be exploited by SCHNAPPs including cell-specific cytometry data from MARS-seq (Jaitin et al., 2014), chromatin accessibility data from ATAC-seq (Buenrostro et al., 2015), or other single-cell approaches. It is then possible to visualize cytometry results from MARS-Seq experiments together with gene expression data from sequencing. Different samples are tracked analogously. The processed data can be exported as singleCellExperiment objects. This allows creating a SingleCellExperiment object from a CSV file that is useable with iSEE or other tools based on the SingleCellExperiment object. Multiple SingleCellExperiment files can be loaded and analyzed together.

**Reproducibility** is achieved by creation of a compressed directory that holds an R-markdown file (Allaire JJ et al. 2019) with associated data archiving all major data manipulations (removal of data, normalization, clustering) and plots that can be saved on request. Thus, a bioinformatician can reproduce the cell selection, validate the analysis steps and optimize the graph for final publication.

All user modifiable parameters in SCHNAPPs can be set individually before starting the application. This is useful when sharing sessions. It is similar to bookmarking with the advantage that no data directory holding the server-side settings has to be shared.

The **shiny framework** (Chang et al., 2019) is used as the underlying framework with the dashboard design (Chang and Borges Ribeiro, 2018) for the graphical user interface (GUI). It can be run from within RStudio (RStudio Team, 2016) or as a stand-alone web application.

The function **schnappsLite** allows publishing precomputed results using the shiny sever. Here, the compute intensive components (normalization, clustering …) have been removed and the number of cells can also be limited. Thus, publication results can be easily made available to the general public. See http://hub05.hosting.pasteur.fr/scProjects/ for examples.

**Internally**, the SingleCellExperiment object is used to store count matrices and user-supplied annotation. To take full advantage of the reactive concept with its dependency graph (Chang et al., 2019), individual computations (normalizations, projections/covariates) are stored in distinct objects. This approach avoids recalculating objects that do not depend on parameters that have changed. Due to the low coverage and dropouts associated with most single cell sequencing experiments, sparse matrices are used to represent raw and normalized read counts, reducing the memory footprint. Parallel implementations are used when available, such as for tSNE (Donaldson, 2016), UMAP (Melville, 2019), and self-organizing maps (SOM; Wittek et al., 2017; can be found in the contributions). Shiny modules allow reuse and standardization of visualizations. Violin plots, 2D plots, and tables are modularized and can be used by any other contributed functionality. The 2D plot module, for example, allows to select cells, re/define/name groups of cells, log transform data, or to normalize it by e.g. gene count per cell, just for the given plot. Selected cell names can optionally be shown and thus copied and pasted. This provides the ability to refine the analysis by e.g. sub-clustering a set of cells within a given cluster and represents an important tool for the bench-scientist who is looking to identify the phenotype and potential fate of cells.

**Contributions** allow adding analyses or visualization tools; the end-user provides the directory where the contributions are located on the file system during startup of the application. The application then looks for specific file names that contain sources for the GUI elements and reactive objects. Contributions for trajectory inference (SCORPIUS (Cannoodt R. et al., 2016), ElPiGraph.R (Albergante, 2019)), for imputation (DCA (Eraslan G et al., 2019), and SOMs are already available at github.com/baj12/SCHNAPPsContributions. A dummy contribution is available that holds example code for key features (adding projections, normalizations/imputations, visualizations, and reports), which allows developers an easy entry point. New normalization and imputation methods can be integrated as well as differential expression methods. This concept allows reducing the functionality to only those tools that are useful for a given biological question and thus reduce the complexity of the application.

A key concept within SCHNAPPs is that all single attributes of a cell (other than the count for a given gene) are treated equally. That means that we don’t distinguish between cluster assignment and projections as both are numerical (or factorial) vectors and we call them all projections (they allow plotting the high dimensional data into lower (2/3) dimensions). Cell-type assignments are added as new projections as well as time assignments coming from a trajectory inference.

A **computer** with substantial memory and CPUs is recommended for the use of SCHNAPPs (e.g. 64GB RAM allows working with ~40,000 cells and ~15,000 genes), though the supplied example data (200 cells, 800 genes) should be loadable on any modern computer.

## Example use case and discussion

**A typical workflow** on how to use the application is described in the following, showing a non-exhaustive list of key features. An in-depth walkthrough and comparison to a scran workflow (https://bioconductor.org/packages/release/bioc/vignettes/scran/inst/doc/scran.html) is given in the supplements.

1. Prepare input data by creating a SingleCellExperiment object in R with all available metadata and save this in an “RData” file. This step is best performed by a trained bioinformatician who can prefilter and precompute different properties of the data set. The supplementary also gives an example on how to setup the data.

2. Lanuch SCHNAPPs and load the input data.

3. Check the quality of the data by looking at the unique molecular identifiers (UMIs) (*General QC - UMI histogram, figure 1A*), the distribution of samples (*General QC - Sample histogram, figure1,B*), and the distribution of highly expressed genes (*General QC - Scater QC*, figure1 C).

**Figure 1.**
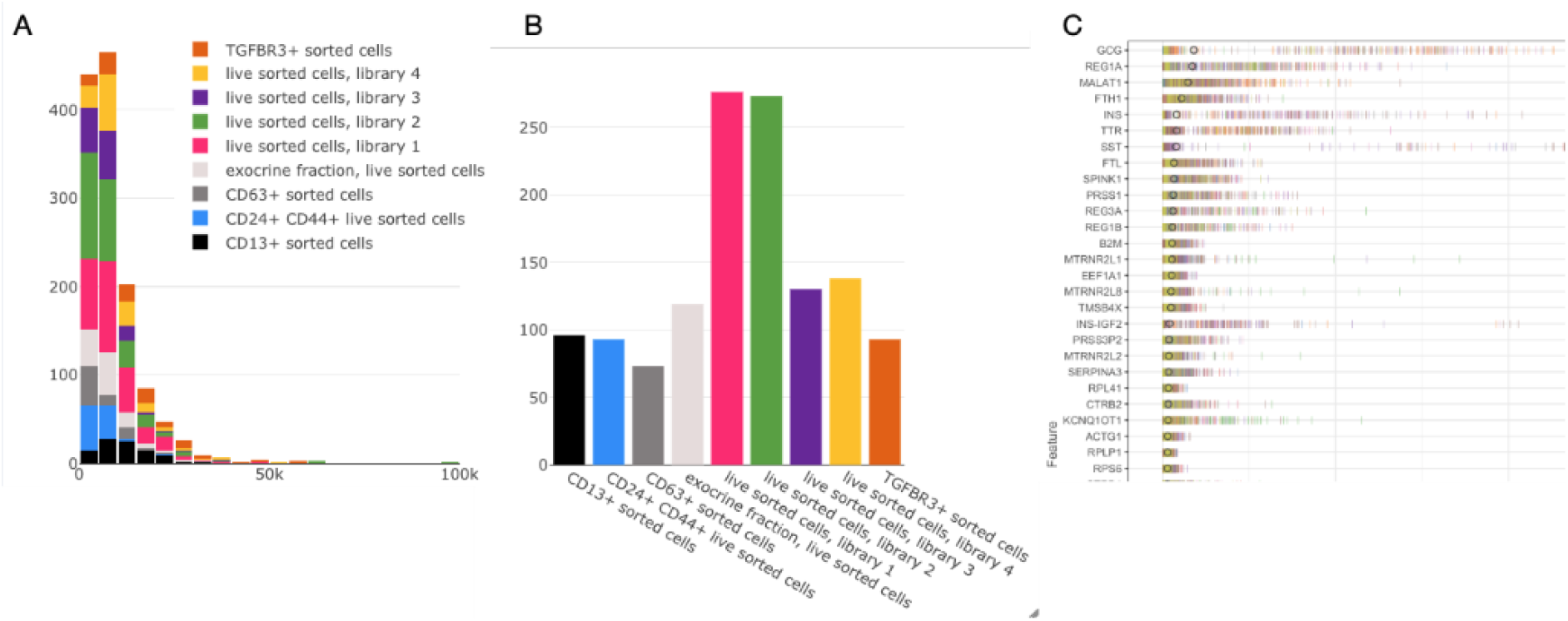
Selection of quality control figures. Data shown is from the workflow that follows the scran data analysis vignette (Supplementary document) with data from (Grun D. et al., 2016). **A**: histogram of UMIs per cell and sample (indicated by color). The number of bins can be chosen by the user, zooming is available to identify a threshold for min/max expression per cell. **B**: histogram of cells per sample. **C**: distribution of reads per cell for highly expressing genes (y-axis). Individual dashes represent the abundance of a given gene for an individual cell colored by sample. X-axis shows the number of UMIs.

4. Using the 2D plot under “*Co-expression - selected*” identify cells that should be removed (figure 2, A). Copy the cell names that are shown after checking “*show more options*” followed by checking “*show cell names*” (figure 2, B).

**Figure 2.**
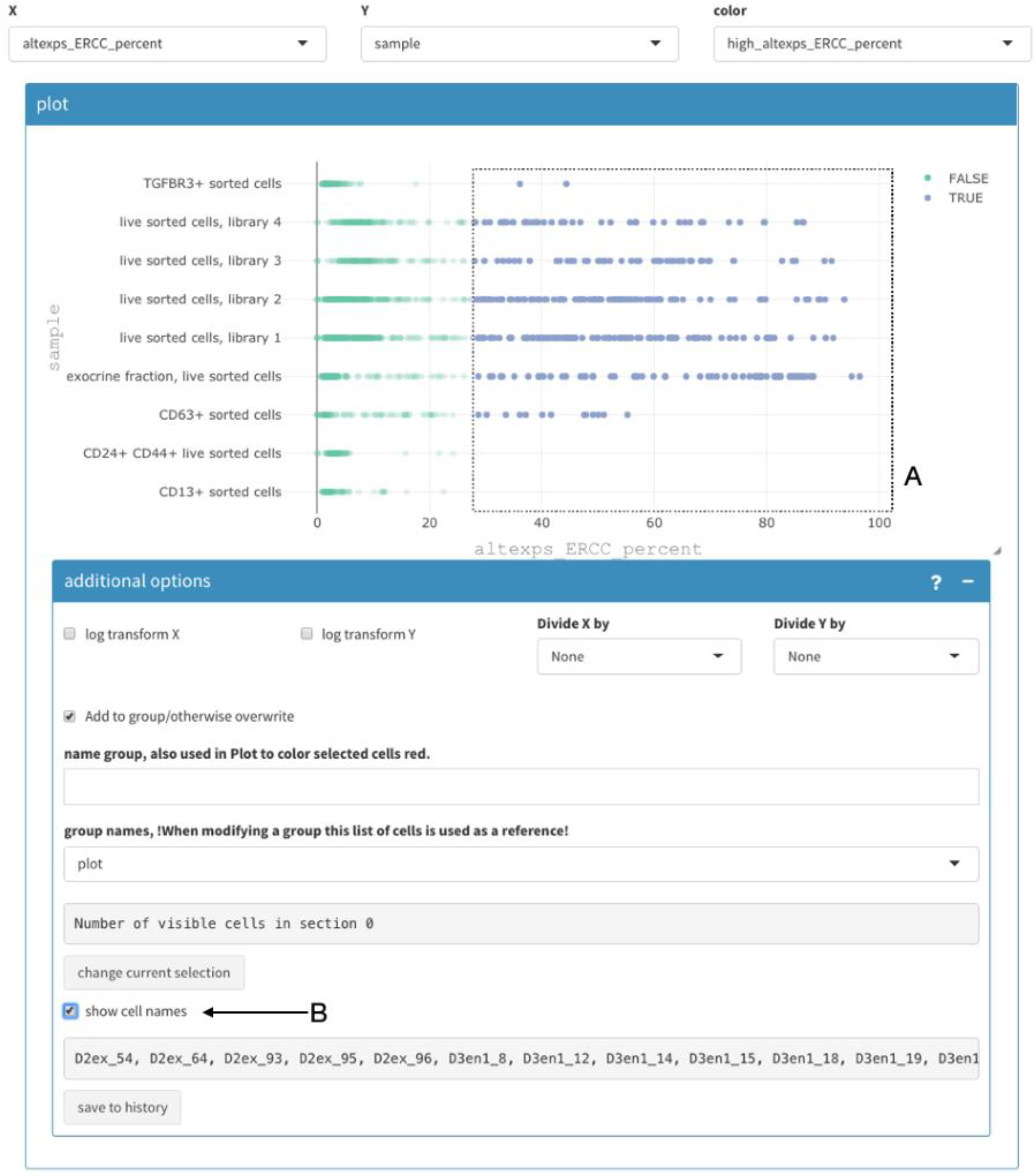
Selecting cells in 2D plot. The precalculated “altexps_ERCC_percent” (X-axis) is used to select cells that should be removed. **A**: Manual selection of cells. **B**: Show cell names to copy and paste in the appropriate field to remove cells. See supplementary document 1 for more details.

**Figure 3.**
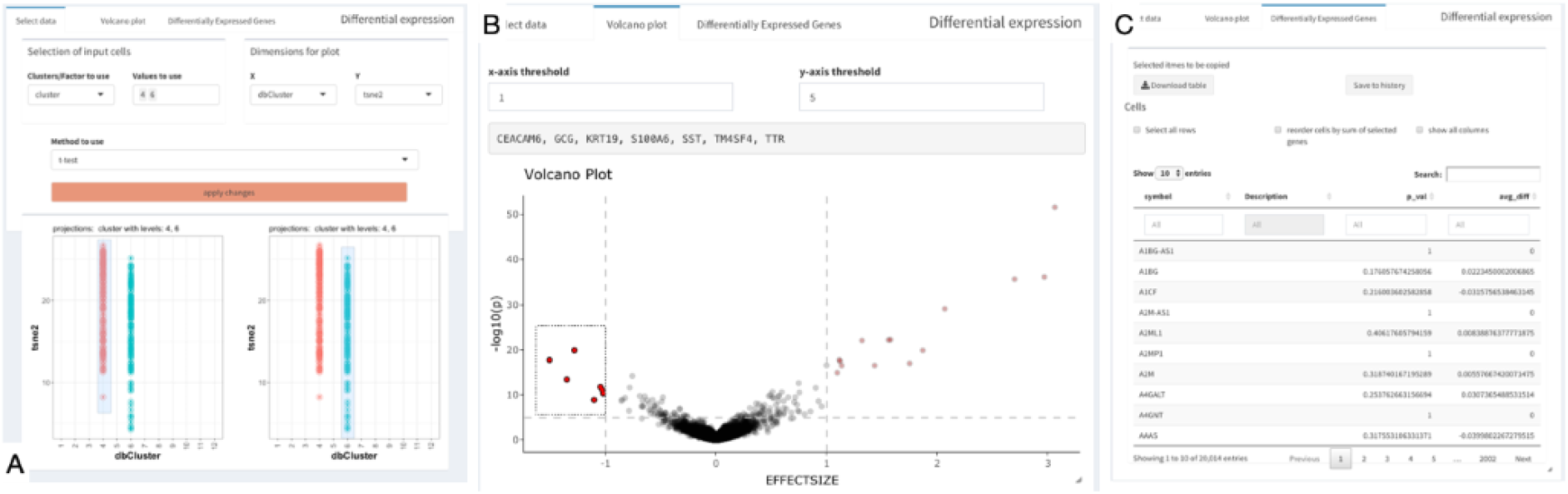
Differential gene expression analysis. **A**: Selection of cells that should be compared. It is possible to pre-filter cells. Here, only cells from clusters 4 and 6 are shown. The t-test is selected as method to use. **B**: resulting Volcano plot. The selected genes are named above the plot and copied from there. **C**: table with differentially expressed genes and key analysis and descriptive values.

5. Paste this selection under “*Cell selection - Cells to be removed*”, alternatively select cells that are pertinent for the question at hand (*Cell selection - cells to keep (remove others)*) then click “apply changes” in Cell selection to update the underlying data. This will trigger recalculating all projections and renormalizing of the data. It is also possible to apply thresholds for minimal/maximal number of UMIs, require that certain genes are either expressed or not. Genes can be removed using regular expressions for the gene symbol or minimal number UMIs over all cells.

Eight different normalization methods are implemented. Parameters for principle component analysis (PCA) can be changed, including the number of variable genes to be used and how to calculate these highly variable genes, among others. Different cluster methods are available based on the standard workflows from Seurat (https://satijalab.org/seurat/vignettes.html) and scran.

6. Verify that the number of cells is correct by looking at the summary statistics on the sidebar. These summary statistics list the number of cells before and after normalization (certain normalization methods remove genes), the file names being used, and the median number of UMIs and genes.

7. Repeat 4-6 until satisfied with the cell selection.

8. Identify genes of interest by performing a comparative analysis between two groups of cells (*Subcluster analysis - DGE analysis*).

9. Investigate how different genes are co-expressed using any of the following: *Co-expression - All cluster* shows in a heat-map clusters/samples of a given set of genes, or the genes that distinguish between clusters (findMarkers from the scran package) if the gene list is empty; *Co-expression - Violin plot*, which visualizes the number of cells expressing a set of genes or any combination of them (Figure 4 A); *Co-expression - SOM cluster*, which calculates a SOM and shows genes that co-cluster with a gene of interest; *Data Exploration - Panel plot*, which shows 2D projections for a given set of genes.

**Figure 4.**
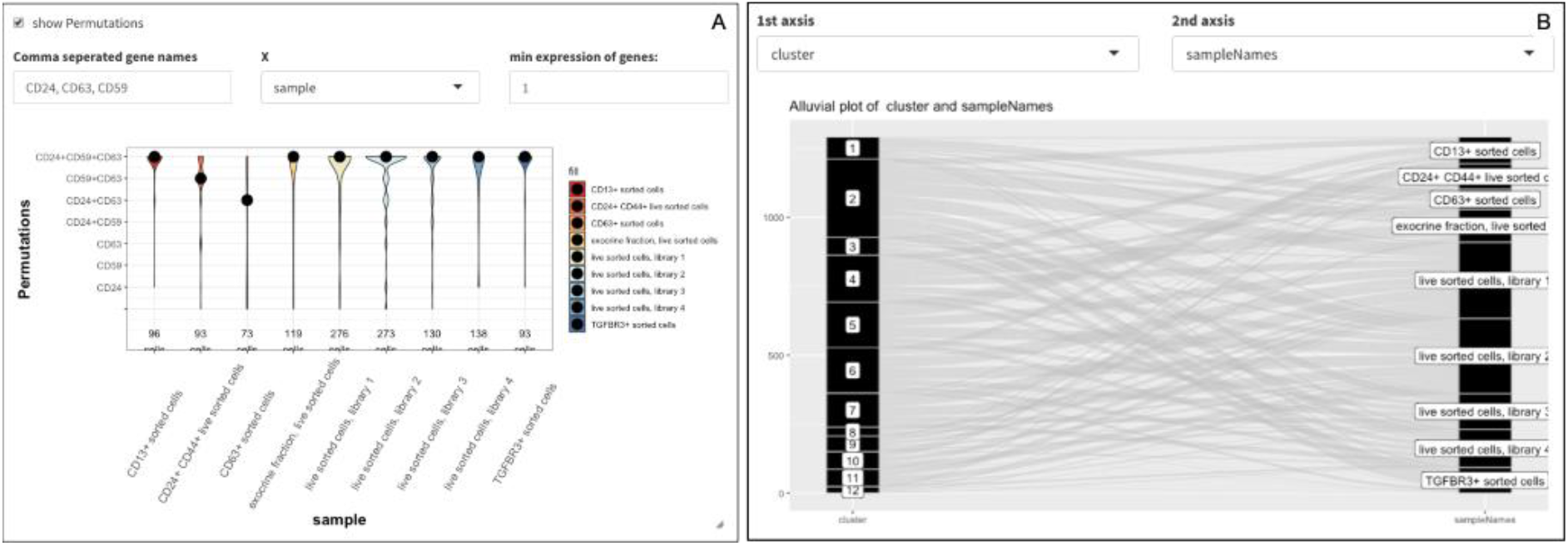
Two examples of plots that can be generated using SCHNAPPs. A: Violin plot showing the co-expression of three genes for each sample. B: Alluvial plot between cluster association and samples.

10. Select cells based on criteria like the expression of a given gene (*Co-expression - selected - more options - group name*) and name the selection (this creates a new projection that can be used elsewhere).

11. Export data as RData file with all parameters used. This can also be used in the SCHNAPPs-lite version enabling faster access to the data with less compute resources.

12. Download the history of associated data and HTML report as a compressed directory as described earlier. Transmit this to a bioinformatician for validation and optimization.

13. It is also possible to bookmark a given state of the application, and thus come back to a given state of the analysis.

Colors used for samples and clusters can be changed. Projections can be combined, renamed, and factors can be releveled. This allows creating and annotating groups of cells and comparing different analysis strategies. Basic analyses of **cell populations** can be performed using histograms, violin plots, or heatmaps. Nine different methods for differential gene expression analysis are implemented.

This gives just a small glimpse of the possibilities. The 2D plot alone, with its potential to show any combination of projections, meta-data, and groupings, allows for many quality control and cell selection opportunities. To better guide the user there are several vignettes, FAQs and other information on GitHub (https://c3bi-pasteur-fr.github.io/UTechSCB-SCHNAPPs/articles/pkdown/SCHNAPPs_usage.html#co-expression).

SCHNAPPs is an established (Neri T, et al, 2019; Suffee N, et al., 2020) analysis and visualization tools for a) bench-scientists to autonomously explore and communicate his data and b) bioinformaticians to validate and continue this work and easily integrate new functionality.

## Supporting information

Supplemental Workflow

## Acknowledgements

We would like to thank the members of Single cell working group Pasteur/Paris for helpful discussions; Anna Barcons, Eric Tartour, Antonin Saldmann, Mandar Patgaonkar, Lisa Chakrabarti, and James DiSanto for testing and working with scShinyHub and SCHNAPPs. Valentina Libri for testing, reading the manuscript, and helpful discussions. Kenneth Smith for careful reading of the manuscript.

