## Supplemental Workflow for "SCHNAPPs - Single Cell sHiNy APPlication(s)": FollowScranWorkflowToAnalyzeSingleCellRNAseqData.html

Follow scran workflow to analyze single-cell RNA-seq data


Code 

- Show All Code
- Hide All Code

### Follow scran workflow to analyze single-cell RNA-seq data

###### Bernd Jagla

bernd  jagla  pasteur  fr


#### 2.6.2020

```
library(scran)
library(scater)
library(SCHNAPPs)
library(BiocParallel)
register(SerialParam()) # avoid problems with fastMNN parallelization.
set.seed(100)
```

### Introduction

This vignette shows how SCHNAPPs can be applied as a single-cell RNA-sequencing platform by comparing it to a standard workflow as decribed under SCRAN (https://bioconductor.org/packages/release/bioc/vignettes/scran/inst/doc/scran.html).

### Data setup

We pre-compute the scran workflow and make its results available for comparison within SCHNAPPs.

We use the same data set. Here, we have to create a singleCellExperiment object and save it in an external file to be able to load into SCHNAPPs later.

```
# BiocManager::install("scRNAseq")
library(scRNAseq)
sce <- GrunPancreasData()
sce
## class: SingleCellExperiment 
## dim: 20064 1728 
## metadata(0):
## assays(1): counts
## rownames(20064): A1BG-AS1__chr19 A1BG__chr19 ... ZZEF1__chr17
##   ZZZ3__chr1
## rowData names(2): symbol chr
## colnames(1728): D2ex_1 D2ex_2 ... D17TGFB_95 D17TGFB_96
## colData names(2): donor sample
## reducedDimNames(0):
## spikeNames(0):
## altExpNames(1): ERCC
```

We prepare the singleCellExperiment object to contain the col/row Data that is needed by SCHNAPPs. At this point, which is usually performed by the bioinformatician who is preparing the data, we can also add other information and documentation.

colData has to contain the columns barcode = a unique identifier per experiment sampleNames = a name for a collection of cells.

rowData has to contian the columns symbol = the gene symbol Description = a text describing the gene id = unique identifier, e.g. ENSG number

```
qcstats <- scater::perCellQCMetrics(sce)
colData(sce) = cbind(colData(sce), qcstats)
colData(sce)$barcode = rownames(colData(sce))
colData(sce)$sampleNames = factor(colData(sce)$sample)

rowData(sce)$Description = ""
rowData(sce)$id = rownames(rowData(sce))
```

To show the effect of the scater filtering we also calculate those. Missing values are handled as genes with high ERCC expressing genes.

```
qcfilter <- quickPerCellQC(qcstats, percent_subsets="altexps_ERCC_percent")
qcfilter[is.na(qcfilter$high_altexps_ERCC_percent),"high_altexps_ERCC_percent"] = T
colData(sce) = cbind(colData(sce), qcfilter)
```

```
# here we redo the scran workflow with different clustering algorithms 
# and store the results in the singleCellExperiment object that we are going to use in SCHNAPPs
sceOrg = sce
sce <- sce[,!qcfilter$discard]
# summary(qcfilter$discard)

library(scran)
clusters <- quickCluster(sce)

sce <- computeSumFactors(sce, clusters=clusters)
# summary(sizeFactors(sce))

sce <- logNormCounts(sce)

dec <- modelGeneVar(sce)
# plot(dec$mean, dec$total, xlab="Mean log-expression", ylab="Variance")
# curve(metadata(dec)$trend(x), col="blue", add=TRUE)

dec2 <- modelGeneVarWithSpikes(sce, 'ERCC')

# Get the top 10% of genes.
top.hvgs <- getTopHVGs(dec, prop=0.1)
sce <- runPCA(sce, subset_row=top.hvgs)
# reducedDimNames(sce)
sced <- denoisePCA(sce, dec2, subset.row=getTopHVGs(dec2, prop=0.1))
# ncol(reducedDim(sced, "PCA"))
output <- getClusteredPCs(reducedDim(sce))
npcs <- metadata(output)$chosen
reducedDim(sce, "PCAsub") <- reducedDim(sce, "PCA")[,1:npcs,drop=FALSE]
# npcs
# In this case, using the PCs that we chose from getClusteredPCs().
g1 <- buildSNNGraph(sce, use.dimred="PCAsub")
cluster <- igraph::cluster_walktrap(g1)$membership
sce$cluster <- factor(cluster)
# table(sce$cluster)

g1a <- buildSNNGraph(sce, use.dimred="PCAsub")
cluster <- igraph::cluster_walktrap(g1a)$membership
sce$cluster1a <- factor(cluster)

# In this case, using the PCs that we chose from getClusteredPCs().
g <- buildSNNGraph(sce, use.dimred="PCAsub", type = "jaccard")
cluster <- igraph::cluster_walktrap(g)$membership
sce$cluster2 <- factor(cluster)

g <- buildSNNGraph(sce, use.dimred="PCAsub", type = "number")
cluster <- igraph::cluster_walktrap(g)$membership
sce$cluster3 <- factor(cluster)


# table(sce$cluster) 
sce <- runTSNE(sce, dimred="PCAsub")
# plotTSNE(sce, colour_by="cluster", text_by="cluster")
# plotTSNE(sce, colour_by="cluster2", text_by="cluster2")
# plotTSNE(sce, colour_by="cluster3", text_by="cluster3")


library(pheatmap)
ass.prob <- bootstrapCluster(sce, FUN=function(x) {
    g <- buildSNNGraph(x, use.dimred="PCAsub")
    igraph::cluster_walktrap(g)$membership
}, clusters=sce$cluster)

sceOrg$cluster = 0
sceOrg$cluster1a = 0
sceOrg$cluster2 = 0
sceOrg$cluster3 = 0
names(sceOrg$cluster) = colnames(sceOrg)
names(sceOrg$cluster1a) = colnames(sceOrg)
names(sceOrg$cluster2) = colnames(sceOrg)
names(sceOrg$cluster3) = colnames(sceOrg)
sceOrg$cluster[colnames(sce)] = sce$cluster
sceOrg$cluster1a[colnames(sce)] = sce$cluster1a
sceOrg$cluster2[colnames(sce)] = sce$cluster2
sceOrg$cluster3[colnames(sce)] = sce$cluster3
sceOrg$cluster = as.factor(sceOrg$cluster)
sceOrg$cluster1a = as.factor(sceOrg$cluster1a)
sceOrg$cluster2 = as.factor(sceOrg$cluster2)
sceOrg$cluster3 = as.factor(sceOrg$cluster3)

# pheatmap(ass.prob, cluster_cols=FALSE, cluster_rows=FALSE,
#     col=colorRampPalette(c("white", "blue"))(100))
```

We save all results including the cluster assignments from the scran workflow to be able to compare to SCHNAPPs.

```
save(file = "scran.grunPancreas.RData", list = c("sceOrg"))
```

### SCHNAPPs

Now start SCHNAPPs.

Please see the help and other documentation on how to enable tracking data changes and saving plots (historyPath option). The defaultValues allows us to preset any user defineable variable in the GUI.

This is partially done to simplify your life as you don’t have to set all the parameters partially to avoid mistakes and for this to show the correct set-up.

The variables are described in the code chunk. To note: the number of genes to use is set to 951, which is the same value as in the scran vignette. We don’t have a way to calculate this. But we are aware that changing this value has substaintial effects for the cluster assignment. We strongly believe that the optimization of these parameters is not trivial and we are currently looking for solutions how to solve this. Similar argumenation goes for the number of PCs to use for the clustering algorithm.

```
library(SCHNAPPs)
defaultValues = list()

# input
defaultValues[["sampleInput"]] =FALSE # don't subsample
defaultValues[["whichscLog"]] = "calcLog" # calculate normalization

# Normaliztion
defaultValues[["normalizationRadioButton"]] = "DE_scaterNormalization" # use scaterNorm

# General parameters
## PArameters for PCA
defaultValues[["pcaScale"]] = FALSE # don't scale
defaultValues[["pcaN"]] = 951 # number of genes to use.
defaultValues[["pcaRank"]] = 14 # number PCs to use for clustering


# Parameters for clustering
defaultValues[["tabsetCluster"]] = "snnGraph" # use SNN graph function

# preset Alluvial plot axes
defaultValues[["alluiv1"]] = "cluster"
defaultValues[["alluiv2"]] = "dbCluster"


# Gene selection
# we usually remove genes from mitochrondrium and Ribosome associated proteins using the following regular expression ("^MT-|^RP|^MRP"). This is not done in the scran vignette 
defaultValues[["selectIds"]] = ""
defaultValues[["minGenes"]] = 1 # min number of reads per cell


# Cell selection
defaultValues[["minGenesGS"]] = 1 # minimum number of UMIs (sum) over all cells.
# cells to be removed
defaultValues[["cellsFiltersOut"]] = "D2ex_54, D2ex_64, D2ex_93, D2ex_95, D2ex_96, D3en1_8, D3en1_9, D3en1_12, D3en1_14, D3en1_15, D3en1_18, D3en1_19, D3en1_20, D3en1_21, D3en1_22, D3en1_23, D3en1_25, D3en1_31, D3en1_32, D3en1_34, D3en1_35, D3en1_36, D3en1_39, D3en1_43, D3en1_44, D3en1_47, D3en1_51, D3en1_52, D3en1_53, D3en1_57, D3en1_58, D3en1_59, D3en1_61, D3en1_62, D3en1_64, D3en1_66, D3en1_70, D3en1_72, D3en1_74, D3en1_76, D3en1_77, D3en1_78, D3en1_80, D3en1_82, D3en1_86, D3en1_91, D3en1_92, D3en1_95, D3en1_96, D3en2_1, D3en2_4, D3en2_8, D3en2_9, D3en2_10, D3en2_11, D3en2_14, D3en2_15, D3en2_18, D3en2_19, D3en2_20, D3en2_28, D3en2_29, D3en2_34, D3en2_35, D3en2_36, D3en2_40, D3en2_41, D3en2_43, D3en2_44, D3en2_46, D3en2_47, D3en2_49, D3en2_53, D3en2_54, D3en2_59, D3en2_65, D3en2_67, D3en2_69, D3en2_70, D3en2_75, D3en2_76, D3en2_77, D3en2_78, D3en2_79, D3en2_80, D3en2_82, D3en2_84, D3en2_86, D3en2_87, D3en2_88, D3en2_89, D3en2_95, D3en2_96, D3en3_3, D3en3_4, D3en3_12, D3en3_13, D3en3_14, D3en3_15, D3en3_17, D3en3_19, D3en3_20, D3en3_22, D3en3_23, D3en3_24, D3en3_25, D3en3_26, D3en3_31, D3en3_33, D3en3_35, D3en3_37, D3en3_38, D3en3_42, D3en3_43, D3en3_45, D3en3_47, D3en3_49, D3en3_56, D3en3_57, D3en3_58, D3en3_60, D3en3_61, D3en3_62, D3en3_63, D3en3_66, D3en3_69, D3en3_70, D3en3_71, D3en3_76, D3en3_79, D3en3_88, D3en3_90, D3en3_91, D3en3_93, D3en3_96, D3en4_1, D3en4_5, D3en4_10, D3en4_11, D3en4_13, D3en4_15, D3en4_17, D3en4_18, D3en4_21, D3en4_24, D3en4_27, D3en4_28, D3en4_29, D3en4_34, D3en4_36, D3en4_37, D3en4_40, D3en4_44, D3en4_45, D3en4_49, D3en4_50, D3en4_51, D3en4_52, D3en4_53, D3en4_55, D3en4_57, D3en4_59, D3en4_62, D3en4_66, D3en4_67, D3en4_70, D3en4_75, D3en4_76, D3en4_77, D3en4_78, D3en4_79, D3en4_80, D3en4_88, D3en4_93, D3en4_94, D3en4_95, D3ex_1, D3ex_2, D3ex_7, D3ex_8, D3ex_10, D3ex_11, D3ex_13, D3ex_14, D3ex_15, D3ex_17, D3ex_18, D3ex_19, D3ex_20, D3ex_21, D3ex_22, D3ex_23, D3ex_25, D3ex_26, D3ex_27, D3ex_28, D3ex_29, D3ex_30, D3ex_31, D3ex_32, D3ex_33, D3ex_35, D3ex_37, D3ex_38, D3ex_42, D3ex_43, D3ex_44, D3ex_45, D3ex_46, D3ex_47, D3ex_49, D3ex_50, D3ex_52, D3ex_53, D3ex_56, D3ex_57, D3ex_58, D3ex_59, D3ex_60, D3ex_61, D3ex_62, D3ex_63, D3ex_64, D3ex_65, D3ex_67, D3ex_69, D3ex_70, D3ex_72, D3ex_73, D3ex_75, D3ex_78, D3ex_79, D3ex_80, D3ex_81, D3ex_82, D3ex_84, D3ex_85, D3ex_86, D3ex_89, D3ex_93, D3ex_94, D3ex_95, D3ex_96, D74_4, D74_11, D74_19, D74_30, D74_38, D74_50, D74_72, D74_80, D71_13, D71_24, D71_25, D71_46, D71_50, D71_74, D71_77, D71_96, D72_6, D72_17, D72_26, D72_51, D72_56, D72_60, D73_53, D73_66, D73_74, D73_75, D73_77, D73_80, D73_88, D10631_1, D10631_6, D10631_13, D10631_16, D10631_28, D10631_30, D10631_31, D10631_32, D10631_35, D10631_49, D10631_59, D10631_77, D10631_86, D101_2, D101_3, D101_4, D101_6, D101_9, D101_12, D101_15, D101_18, D101_19, D101_20, D101_23, D101_26, D101_30, D101_32, D101_33, D101_38, D101_39, D101_40, D101_41, D101_47, D101_49, D101_54, D101_55, D101_60, D101_61, D101_62, D101_63, D101_64, D101_65, D101_67, D101_68, D101_70, D101_71, D101_75, D101_76, D101_78, D101_79, D101_80, D101_85, D101_86, D101_87, D101_90, D101_92, D101_94, D101_96, D102_2, D102_5, D102_8, D102_9, D102_10, D102_12, D102_13, D102_15, D102_16, D102_19, D102_20, D102_22, D102_23, D102_27, D102_28, D102_29, D102_30, D102_31, D102_32, D102_33, D102_34, D102_35, D102_36, D102_37, D102_38, D102_41, D102_42, D102_43, D102_44, D102_46, D102_48, D102_49, D102_52, D102_53, D102_56, D102_60, D102_62, D102_65, D102_67, D102_70, D102_72, D102_74, D102_77, D102_84, D102_85, D102_88, D102_89, D102_90, D102_92, D102_95, D17All1_13, D17All1_94, D17All2_5, D17All2_50, D17All2_89, D17TGFB_60, D17TGFB_62, D2ex_94, D3en1_40, D3en2_51, D3en3_18, D3en3_40, D3en3_77, D3en3_81, D3en3_83, D3en4_90, D3en4_96, D74_48, D74_84, D74_96, D71_89, D72_94, D73_47, D10631_47, D10631_75, D10631_78, D10631_79, D10631_80, D10631_83, D10631_84, D10631_94, D101_8, D101_11, D101_46, D102_47, D102_80, D102_81, D102_83, D172444_94, D172444_95, D172444_96, D17All1_14, D17All1_15, D17All1_17, D17All1_85, D17All2_96, D17TGFB_96"

schnapps(defaultValues = defaultValues)
```

#### Removal of cells

This section shows how to remove the cells that have been identified in the scran vignette. To follow we have to remove the cells already selected. and set “Cells to be removed” under Cell selection - additional parameters to an empty string as shown below:

Filter based on ERCC percentage

and click on apply changes.

Then load the data we just created.

Filter based on ERCC percentage

We now show which filters have been applied to the data in the scran workflow. We use the perCellQCMetrics metrics that cannot be calculated from within SCHNAPPs but could be precalculated. We can then visualize the cells that are filtered out and recover the thresholds used in individual steps:

##### based on ERCC percentage:

This percentage is precalculated from the scr

The data can be visualized in the 2D plot (co-expression - selected) with the following paramters: X : altexps\_ERCC\_percent Y: sample color: high\_altexps\_ERCC\_Percent

Filter based on ERCC percentage

The selected cell names can be copy/pasted to the “Cells to be removed” under Cell selection - additional parameters" as described above.

##### filter low expressing cells

Using a similar 2D plot with the following paramters: X : detected Y: sample color: low\_n\_features

we can select and remove cells with low expressing features as described above.

Filter based on detected genes

We end up with the same 1219 cells as in the vignette from scran.

### Normalization / clustering

We use “scaterNorm”, which is applying exactly the same procedure that is used in the vignette. The user has several other options available that are better described in the online documentation.

The snnGraph method is used for the clustering.

### Results are the same

The alluvial plot between the cluster assignment from the scran package and the cluster assignment from SCHNAPPs (dbCluster) shows that they are the same.

Cluster assignment comparison using Alluvial plot

The same result can be concluded from plotting the cluster assignments in a 2D plot:

Cluster assignment comparison using 2D plot

### Characterization of results

The following will show a few possible results that can be achieved using SCHNAPPs:

#### Co-expression - All clusters

Co-expression - All clusters; After list of gene names has been set to empty SCHNAPPs will automatically fill the list with characteristic genes for each cluster.

The following video shows how to use the heatmap module. If the field with the gene names is empty the the function FindMarkers is used to identify the genes that are most characteristic for each cluster. This list of genes is used for the heatmap.

The genes CPA1, CTRB1, PLA2G1B, PRSS1, PRSS3P2 are highly expressed in cluster 2. GCG, TTR are highly expressed in cluster 5.

#### Co-expression - selected

In the following we describe a few possibilities of the 2D plot

##### co-expression of two lists of genes

We can now compare the genes previously found and see how these cells express genes from different clusters.

2D plot of the sum of expression of two different lists of genes, colored by cluster association.

The cell density can be used to identify where the majority of cells lies or if and where many cells are drawn on top of eachother.

2D plot of the sum of expression of two different lists of genes, colored by density of cells.

The composition of clusters relative to the donor information can be also using the same tool as shown below:

Histogram of cell count per cluster assignment and split by donor information.

UMAP projection. Selecting and naming of a group of cells.

Cells can be grouped and named using the 2D plot, Those are are binary groups, i.e. a cells belongs either to the group or not.

The following video shows how to name groups of cells: \* Cells have to be selected in the 2D plot. \* Give a name \* ensure that the reference for the selection is set to “plot” \* click “change current selection”

To note: \* in-between, the cells seem not to be selected anymore, but that is a “feature” of the underlying visualization functions. \* using the a previously defined group of cells allows to avoid selecting cells that are potentially hidden. This can happen, when the 2D coordinates have changed. This also allows a more cytometry way of thinking, where concurrent selections of cells are used to filter.

After having selected multiple groups, we have these groups as projections available with TRUE/FALSe assignment for each cell. Sometimes, we want to combine these projections, e.g. if there are more than two cell types. This is done under Parameters - Projections - combine projections:

- Select the two projections that should be combined
- name the new projection
- repeat as needed.

Since it is not ensured that these projections are not overlapping, the individual values are concatenated and a new factor is created. The levels of these factors can be changed under the “rename levels” tab. The tables allows verifying the meaning of the levels.

When renaming the levels, don’t forget to add a meaningfull name, otherwise “X” is chosen for you.

If this happens, don’t worry, this can be changed in the “rename projections” tab, where projections can be deleted, but only those you have created manually.

#### Violin plots

The violin plot can show all possible combinations if “show Permutations” is selected, otherwise a normal violoin plot is shown (not shown here).

Violin plot with permutataions

Under “grouped” a second variable can be used. Here, grp1-7 are clusters of cells that have been identified from the UMAP presentation.

grouped violin plots

#### Three dimensional plot

The sum of expression for a list of genes is shown in the 3D tSNE plot that can be found under Data Exploration -

3D tSNE plot with expression of specified genes

Panel Plot

#### Differential gene expression (DGE)

This short video shows how to perform a DGE analysis. The individual steps are also explained afterwards in more detail.

##### selections of groups of cells

The “selection of input cells” allows subsetting the cells even before the visualization.

Cells to be compared can be either selected in the two plot directly, of if cells are only selected in the first plot, all other cells are being used.

DGE between specific groups of cells

##### Volcano plot

Genes can be selected in the volcano plot and their names/symbols are shown above the plot to be copy/pasted.

DGE results as Volcano plot

##### DGE table

A complete table is available, that can also be downloaded.

Table of DGE

#### Comparing different clustering methods

SCHNAPPs uses the projection “dbCluster” for the current clustering result. In order to compare different clustering procedures the rename projection tool allows the user to copy the results do a different name.

Projections can be renamed

### Please help

We hope you find SCHNAPPs useful. Please don’t forget to cite our work when you publish, nor tell us if you find any bugs or have questions/suggestions.

bernd  jagla  pasteur  fr
